# Characterization of transcription activation domain of EcaICE1 and its interaction with EcaSIZ1 in *Eucalyptus camaldulensis*

**DOI:** 10.1101/859819

**Authors:** Ziyang Zhang, Ling Cheng, Weihua Zhang, Jianlin Hu, Yan Liu, Yuanzhen Lin

## Abstract

SUMOylation in plants is associated with biotic and abiotic stress responses, flowering and other aspects of development, and ICE1 protein SUMOylation by SUMO E3 ligase SIZ1 plays important roles in plant cold tolerance. Here, we reported the subcellular localization of EcaICE1 and its interaction with EcaSIZ1 in *Eucalyptus camaldulensis*. The genes *EcaICE1* and *EcaSIZ1* were isolated by homologous cloning. The subcellular localization analysis showed that EcaICE1 was located in nucleus. Bimolecular fluorescence complementation (BiFC) analysis revealed that EcaICE1 could interact with EcaSIZ1 in the nucleus of *Nicotiana benthamiana* leaves. Moreover, yeast two-hybrid assay confirmed that the amino acid region from position 84 to 126 in EcaICE1 was critical for the strong transactivation activity of EcaICE1 and that the C terminal region from position 361 to 557 in EcaICE1 was the key region for its interaction with EcaSIZ1 by using different truncated lengths of non-transactivation activity of EcaICE1 as the bait protein. Collectively, our results showed that EcaICE1 may have a SUMOylation pathway similar to *Arabidopsis thaliana*.

---

Low temperature is one of the important environmental factors that could restrict the growth and development, geographical distribution and production of plants, and low temperature stress could reduce production in agriculture and forest (Janmohammadi et al. 2015). In order to overcome low temperature stress, a series of complex response mechanisms can be activated at physiological and molecular levels for plants (Shi et al. 2018). Researches reveal that a large number of cold-regulated genes (*CORs*) are induced after plant cold acclimation. These *CORs* genes usually contain *DRE*/*CRT* cis-acting elements and can be combined by transcription factors CBFs or DREB1s family (Stockinger et al. 1997). CBFs or DREB1s are key transcription factors of low temperature signaling pathway and play important roles in enhancing plant cold tolerance (Chinnusamy et al. 2007). However, *CBFs* or *DREB1s* are also induced by low temperature. ICE1 gene has been firstly verified from *Arabidopsis thaliana*, which can specifically bind to the cis-acting element of MYC promoter of *CBF3* and then induce the expression of CBF3 downstream gene (Chinnusamy et al. 2003). After that, a lot of studies reveal that ICE-CBF-COR is the important signal network for plants to adapt to the cold stress (Shi et al. 2018). In *Arabidopsis thaliana*, ICE-CBF-COR pathway is positively or negatively controlled by some regulators at transcriptional, post-transcriptional and post-translational levels. SIZ1 (Miura et al. 2007) and OST1 (Ding et al. 2015) are positive regulators, while HOS1 (Dong et al. 2006) and MPK3/6 (Li et al. 2017) are negative regulators. The Ub E3 ligase HOS1, can ubiquitinate and degrade ICE1 protein, thereby increase the cold sensitivity of the transgenic plants (Dong et al. 2006). On the contrary, SUMO E3 ligase SIZ1, can enhance the stability of ICE1 protein by SUMOylation and promote the expression of CBF3, and then causes enhanced cold tolerance (Miura et al. 2007). OST1 can interact with ICE1, competed with HOS1, enhancing the stability and transcriptional activity of ICE1, further resulting in increased cold tolerance (Ding et al. 2015). Recently, the protein kinase MPK3/6 can interact with and phosphorylate ICE1, enhancing the degree of ubiquitination of ICE1, but the interaction between HOS1 and ICE1 is not affected, suggesting that the degradation of ICE1 protein may be performed by the unknown E3 ligase (Li et al. 2017). Although ICE1 gene had been isolated from *A. thaliana* (Chinnusamy et al. 2003), *Brassica juncea* (Wang et al. 2005), *Populus suaveolens* (Lin et al. 2007), wheat (Badawi et al. 2008), banana (ZHAO et al. 2013), *E. camaldulensis* (Lin et al. 2014), *Pyrus ussuriensis* (Huang et al. 2015) and *Hevea brasiliensis* (Deng et al. 2017), the understanding of its positive and negative regulations is still limited in woody plants.

*Eucalyptus* species are the important commercial tree species worldwide. The molecular regulation mechanism of low temperature stress of *Eucalyptus* could have the potential to increase *Eucalyptus* plantation range. *Eucalyptus* CBF genes has been studied (Kayal et al. 2006; Gamboa et al. 2007; Navarro et al. 2009, 2011), but the knowledge about ICE1 or ICE1-like modules is still rather limited. Our precious studies have shown that the overexpression of *EcaICE1* from *E. camaldulensis* can enhance the cold tolerance and the expression level of downstream genes in transgenic tobaccos (Lin et al. 2014). However, the studies of subcellular localization, transcription activity and upstream regulators of EcaICE1 remain uncertain. In this study, we analyzed the subcellular localization and the transcription activation region of EcaICE1. Furthermore, the protein interactions between EcaICE1 and EcaSIZ1 were verified by Bimolecular fluorescence complementation (BiFC) and Yeast two-hybrid (Y2H) assays.

## Methods and Materials

### Plant materials

The 30-day-old rooting tissue culture plantlets of *E. camaldulensis* cv. 103 were used in this study. Tissue culture condition was performed as described previously (Lin et al. 2014). Tobacco (*Nicotiana benthamiana*) plants were maintained in a growth chamber of 25 °C under long day conditions (16/8 h light/dark photoperiod), and 5-week-old plants were selected for further subcellular localization and BiFC analysis.

### RNA isolation and Gene cloning

Total RNA was extracted as described previously (Lin et al. 2014). 1 μg DNase I-treated RNA was used for synthesizing the first strand cDNA according to the manufacturer’s instructions (PrimeScript II 1st Strand cDNA Synthesis Kit; Takara, Dalian, China). The primers of *EcaICE1* and *EcaSIZ1* (listed as table 1) were designed by homology cloning techniques, and then used for amplifying the aim genes with the first strand cDNA as the template. The PCR process as follows: 95 °C for 4 min, followed by 35 cycles of 95 °C for 10 s, 58 °C for 10 s and 72 °C for 20 s.

**Table 1.**
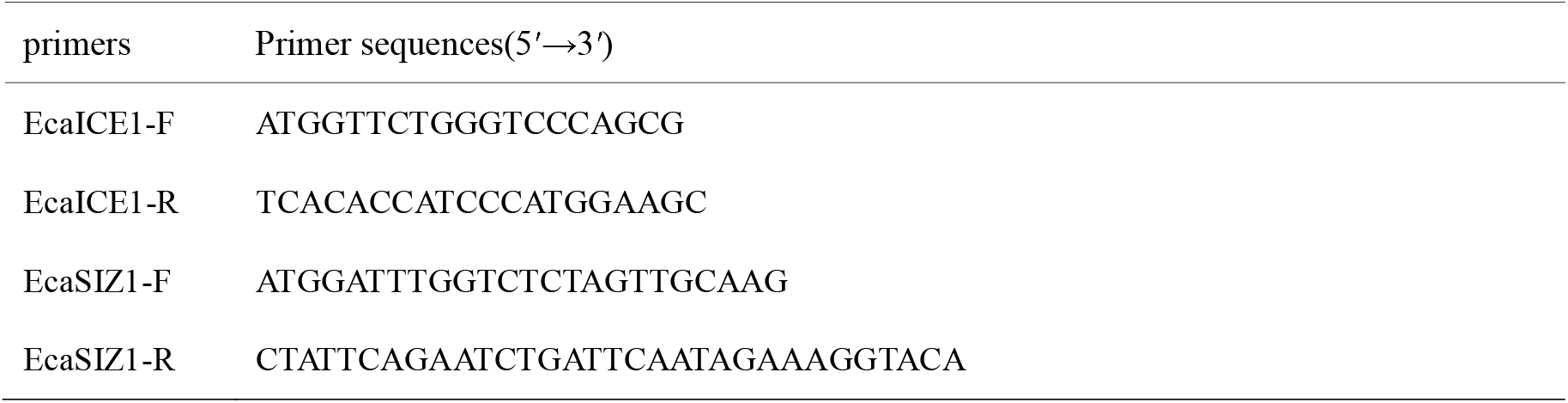
Primers used in the experiments.

### Bioinformatics analysis of gene sequence

The coding sequence (CDS) of *EcaICE1* and *EcaSIZ1* were predicted using FGENESH 2.6 software (http://linux1.softberry.com/berry.phtml?topic=fgenesh&group=programs&subgroup=gfind), and further confirmed by BLASTP program on the NCBI website (http://blast.ncbi.nlm.nih.gov/Blast.cgi). The nuclear localization signals were predicted using ProtComp 9.0 (http://linux1.softberry.com/berry.phtml?topic=protcomppl&group=programs&subgroup=proloc), and the protein secondary domains were searched by Motif scan (https://myhits.isb-sib.ch/cgi-bin/motif_scan). Finally, sequence alignments with other plants were performed with ClUSTALX software.

### Subcellular localization of EcaICE1

The complete Open Reading Frame (ORF) of *EcaICE1* was recombinated to the 5’-terminus of the yellow fluorescent protein (YFP) of the pEarleyGate101 vector with CaMV 35S promoter using primers (listed as table S1). The recombinant vector *35S*∷*EcaICE1*-*YFP* was transferred into *Agrobacterium tumefaciens* strain GV3101, further injected into the abaxial surfaces of 5-week-old tobacco with an incubator for 2-3 d. The YFP signal in the leaves of tobacco was examined using confocal microscopy (Olympus BX61, Tokyo, Japan).

### Bimolecular fluorescence complementation (BiFC) assay

To generate the constructs for BiFC assays, the full-length CDS of *EcaICE1* and *EcaHOS1* (without their stop codons) were subcloned into pUC-pSPYNE or pUCpSPYCE vectors as described previously (Walter *et al*. 2004). Expressions of target genes alone were used as negative controls. The resulting recombinant vectors were used for transient assays of tobacco leaves as described earlier. The transformed tobacco leaves were then incubated at 22 °C for 24–48 h. The YFP signal was observed using a florescence microscope (Zeiss Axioskop 2 plus). The primers for BiFC assay were listed in Supporting Information Table S2.

### Yeast two-hybrid (Y2H) assay

Yeast two-hybrid assays were performed using the Matchmaker™ gold Yeast two-Hybrid Systems (Clontech). Different truncated CDS of *EcaICE1* without transcriptional activation activity and EcaSIZ1 were subcloned into pGBKT7 and pGADT7 vectors to fuse with the DNA-binding domain (DBD) and activation domain (AD), respectively, to create different baits and preys (primers are listed in Supplemental table S3). Then, different pairs of bait and prey recombinants were co-transformed into yeast strain gold Y2H by the lithium acetate method, and yeast cells were grown on SD/-LW medium (minimal media double dropouts, SD medium with-Leu/-Trp) according to the manufacturer’s protocol (Clontech) for 3 days. Transformed colonies were plated onto SD/-LWHA medium (minimal media quadruple dropouts, SD medium with-Leu/-Trp/-Ade/-His) containing 125-μM Aureobasidin A (AbA), to test for possible interactions among EcaICE1 and EcaSIZ1, according to the yeast cell growth status.

### Transcriptional activation assay

The full-length ORF of *EcaICE1* and *EcaSIZ1* were recombinated into pGBKT7 vector, respectively, and co-transformed into yeast strain gold Y2H. Then we found that both EcaICE1 and EcaHOS1 had strong transcriptional activation activity (results are listed in Supplemental Figure S4). So we further performed transcriptional activation assay for EcaICE1. Different truncated coding regions of EcaICE1 (EcaICE1_T1~T6_) were recombinated into pGBKT7 vector, then followed Y2H assay, adding 0, 125, 250, and 500 μM AbA, to further discover the critical region of transactivation activity for EcaICE1. The primers were listed in Supporting Information Table S3.

## RESULT

### The gene amplification and sequencing analysis of EcaICE1 and EcaSIZ1

The multiple alignments of plant ICE1 protein sequences (Figure 1) shows that the *EcaICE1* protein has a highly conserved MYC-like bHLH domain (basic helix-loop-helix) and a zipper structure at the C-terminus, and an S-rich (Serine-rich) acidic domain at the N-terminus. In addition, the SUMO binding site is also present in EcaICE1 and the other two *Eucalyptus* ICE1, and it is identical to ICE1 from *A. thaliana* and *Capsella buras-pastoris*, but slightly different from the other woody plants. Furthermore, the 363-381 position of EcaICE1 protein is a nuclear localization signal, as shown in the NLS(Nuclear localization sequence)box, which means that it may be located in the nucleus. However, it is interesting that only *Eucalyptus* ICE1 proteins have a Q-rich (Glutamine-rich) domain, suggesting that *Eucalyptus* ICE1 proteins might have different characteristics with the other plants.

**Figure 1.**
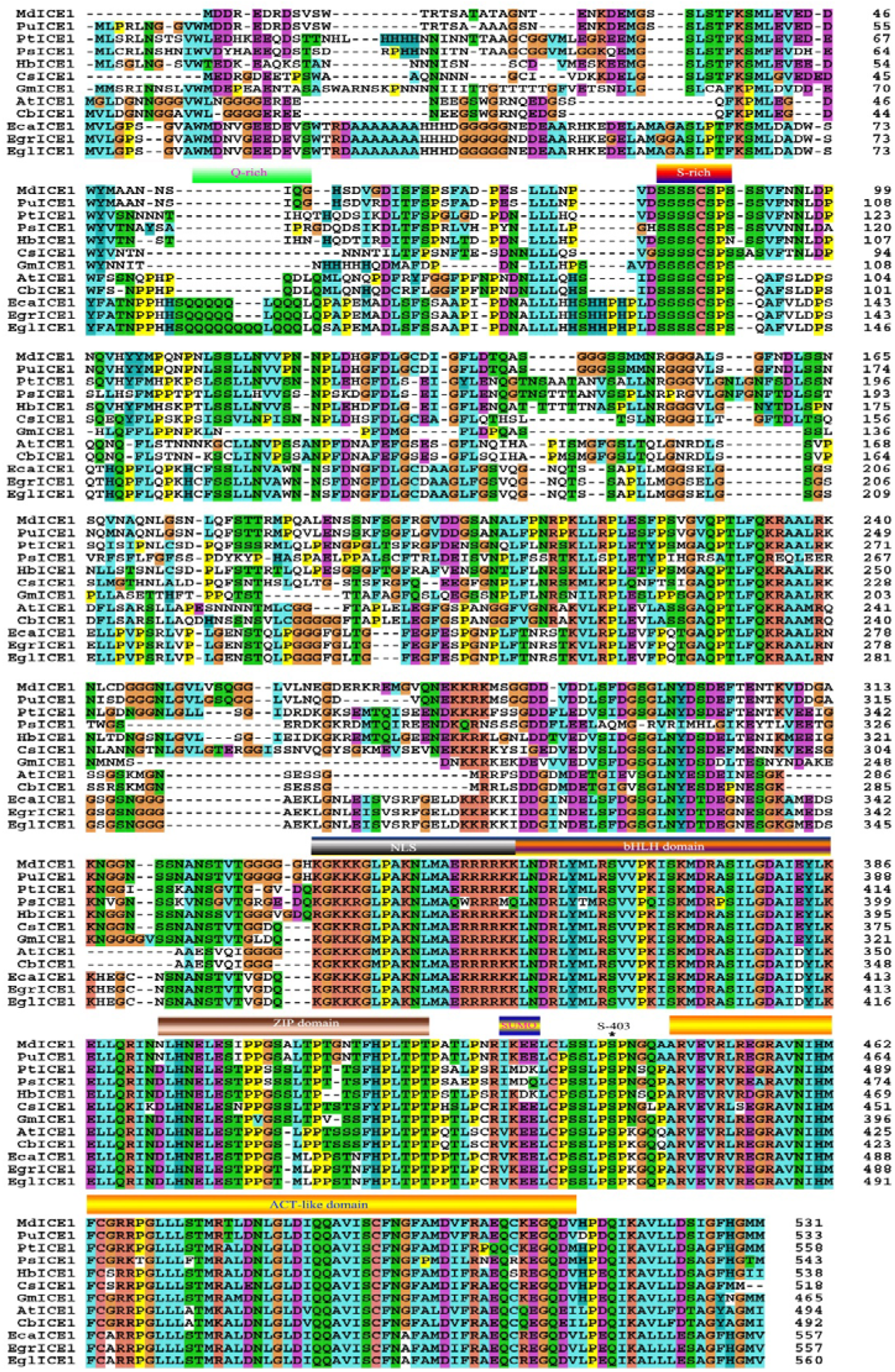
Amino acid alignment of the EcaICE1 protein with twelve plant ICE1 proteins. The predicted protein domains were shown. The *Arabidopsis thaliana* AtICE1 (Accession: AAP14668), *Capsella bursa* CbICE1 (Accession: AAS7935), *Camellia sinensis* CsICE1 (Accession: GQ229032), *Eucalyptus camaldulensis* EcaICE1 (Accession: MH899180), *E. globulus* EglICE1 (Accession: AEF33833), *E. grandis* EgrICE1 (Accession: XP_010066419), *Glycine max* GmICE1 (Accession: ACJ39211), *Hevea brasiliensis* HbICE1 (Accession: JT926796), *Malus domestica* MdICE1 (Accession: ABS50251), *Populus suaveolens* PsICE1 (Accession: ABF48720), *P. trichocarpa* PtICE1 (Accession: ABN58427) and *Pyrus ussuriensis* PuICE1 (Accession: APC57593) proteins are included.

The amplified *EcaSIZE1* cDNA is 2847bp with a full ORF (2616 bp) encoding 872 amino acids. BLAST analysis illustrates that EcaSIZ1 shares a high sequence identity with other plant SIZ1-like proteins, such as *E. grandis* (99%, XP_010043707), *Ziziphus jujube* (72%, XP_015880489), *Malus domestica* (72%, XP_008385287), *Citrus sinensis* (70%, XP_006488140), *P. trichocarpa* (71%, XP_024454498), and *A. thaliana* (68%, OAO90655), respectively. The multiple alignments of plant SIZ1 protein sequences (Figure 2) demonstrate that EcaSIZ1 protein has a conserved MIZ1/SP zinc finger domain, SAP, PINIT, SXS and PHD, similar to other plant HOS1 proteins. These results show that EcaSIZ1 is the SIZ1 protein from *E. camaldulensis*.

**Figure 2.**
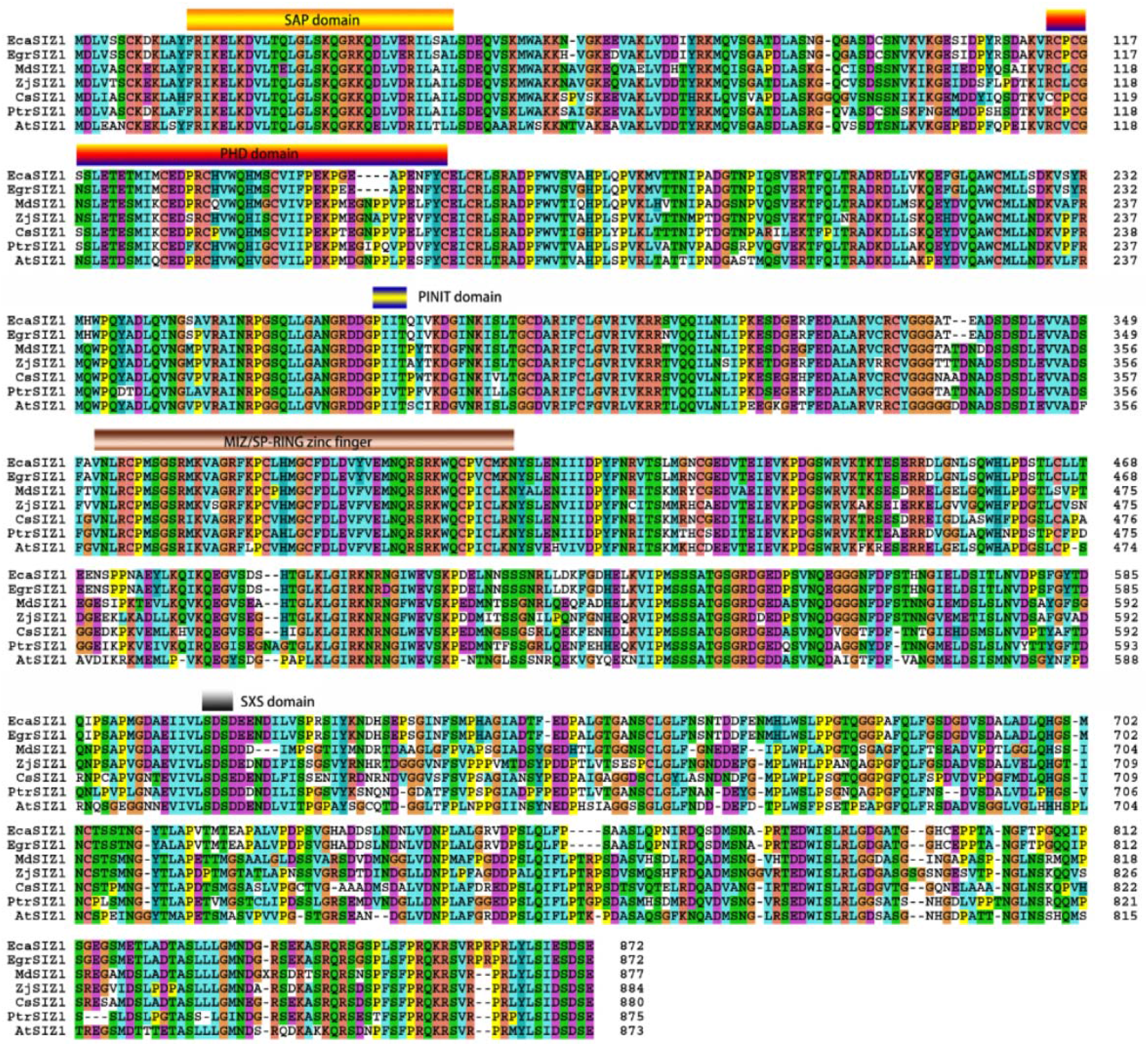
Multi-alignment of amino acid of SIZ1 from *Eucalyptus camaldulensis* and other plants and their conserved domains. *Arabidopsis thaliana*: AtSIZ1, OAO90655; *Citrus sinensis*: CsSIZ1, XP_006488140; *Eucalyptus camaldulensis*: EcaSIZ1, MH899182; *E. grandis*: EgrSIZ1, XP_010043707; *Malus domestica*: MdSIZ1, XP_008385287; *Ziziphus jujuba*: ZjSIZ1, XP_015880489; *Populus trichocarpa*: PtrSIZ1, XP_024454498.

### Subcellular localization of EcaICE1

Based on the multiple alignment analysis (Figure 1), there was one potential NLS sequence in the bHLH domain of EcaICE1. To investigate the subcellular localization of EcaICE1 in plant cells, the full-length coding region of *EcaICE1* was fused to the N–terminus of the YFP reporter gene under the control of the 35S promoter (*35S∷EcaICE1-YFP*). By transient analysis using tobacco leaf epidermis, the yellow fluorescence of EcaICE1 fused-proteins was localized exclusively in the nucleus (Figure 3), indicating that EcaICE1 was a nuclear localized protein, similar to other plant ICE1(Huang et al. 2015; Deng et al. 2017), consistent with the classical role of bHLH as transcriptional regulators in plants (Gabriela et al. 2003).

**Figure 3.**
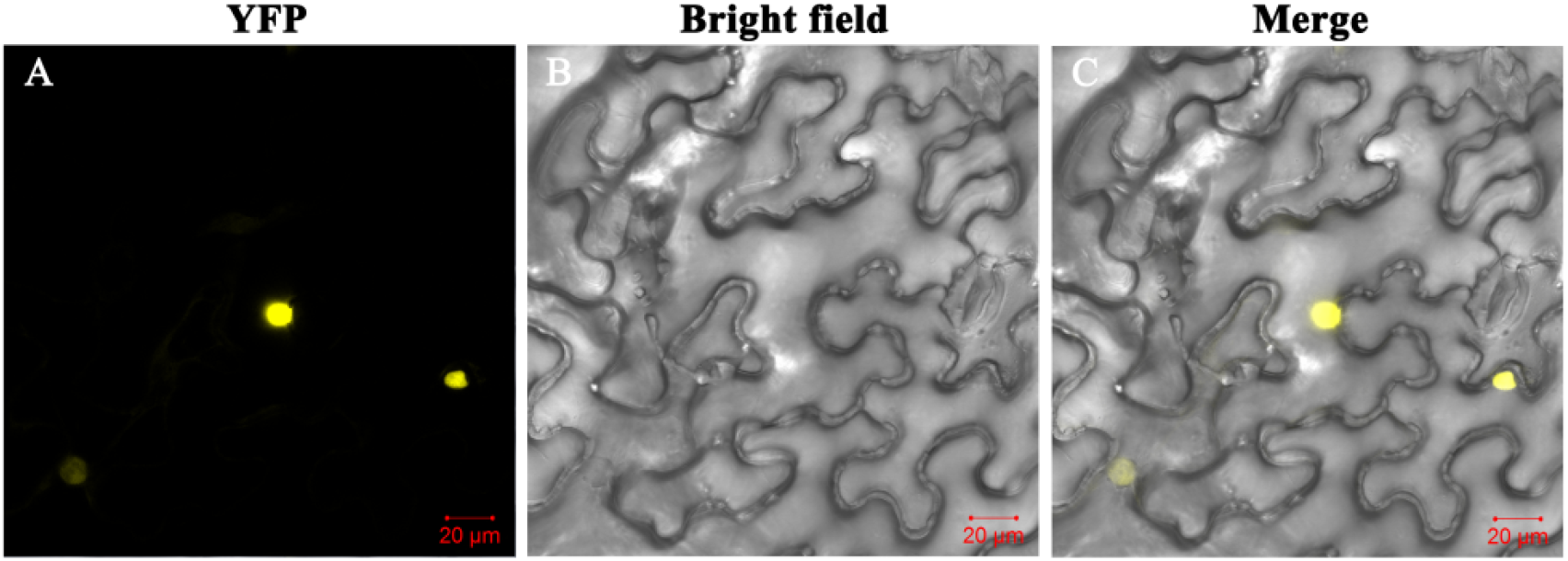
Subcellular localization of EcaICE1 in leaves of *Nicotiana benthamiana*. YFP: the yellow fluorescence signals under YFP-excited; Bright field: figure under bright field; Merge: The merge of YFP and bright field. Bar is 20 μm.

### EcaICE1 could interact with EcaSIZ1

Previous report showed that SIZ1 could interact with ICE1 to regulate its transcriptional activity in *A. thaliana*, which plays an important role in response to cold stress (Miura et al. 2007). In order to investigate whether EcaSIZ1 could also interact with EcaICE1 in *E. camaldulensis*, the protein-protein interactions between EcaICE1 with EcaHOS1 were first analysed using BiFC assay in tobacco leaves. The YFP fluorescent signal was observed at the nucleus when EcaICE1-pSPYCE was co-transfected with EcaHOS1-pSPYNE (Figure 4). In contrast, no YFP fluorescent signal was observed in the negative controls including EcaICE1-pSPYCE co-expressed with unfused pSPYNE or EcaHOS1-pSPYNE co-expressed with unfused pSPYCE. These results showed that EcaICE1 could interact with EcaSIZ1 to form heterodimers at the nucleus (Figure 4).

**Figure 4.**
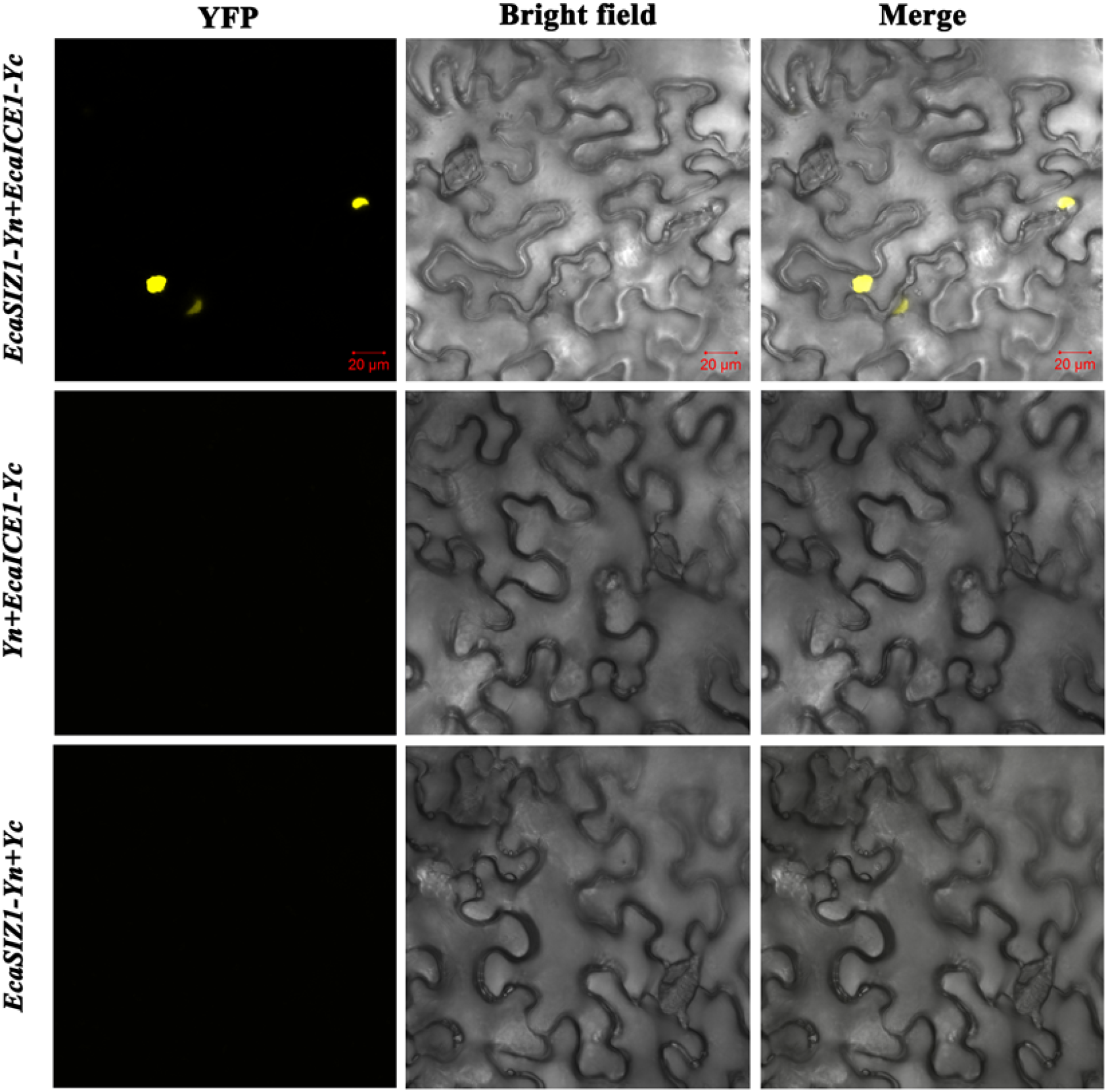
The protein interaction of EcaICE1 and EcaSIZ1 identificated by bimolecular fluorescence complementation methods. YFP: the yellow fluorescence signals under YFP-excited; Bright field: figure under bright field; Merge: The merge of YFP and bright field. Bar is 20 μm.

### Transcriptional activation domain of EcaICE1

To further discover the protein-protein interaction region between EcaICE1 with EcaHOS1, we performed the Y2H assay. It was surprised that not only EcaICE1 but also EcaHOS1 had strong transactivation activity when their full-length sequences were fused into the pGBKT7 vector and co-transfected with pGADT7 empty vector, respectively (Figure S4). In order to remove the effects of strong transactivation activity on protein-protein interaction, we further characterized which region of EcaICE1 acts as transcriptional transactivation, Y2H assays were carried out using intact or truncated EcaICE1 as an effector (Figure 5A). The transfected yeast cells harboring either the full-length EcaICE1 (pGBKT7-EcaICE1) or the truncated version (EcaICE1_T1_(1-41a); EcaICE1_T2_(1-83a)) grew well on the selection medium SD/-LWHA, suggesting that the N-terminal 1-83 residues are not necessary for transactivation activity of the EcaICE1. In contrast, when the N-terminal amino acids of the other truncated version (EcaICE1_T3_(1-125aa); EcaICE1_T4_(1-184aa); EcaICE1_T5_(1-316aa); EcaICE1_T6_(1-360aa)) were deleted, no interactions were detected (Figure 5B). Taken together, these results demonstrate that amino acids from positions 84 to 125 in EcaICE1 are critical for the transactivation activity of EcaICE1, which contains Q-rich domain (Figure 1), in agreement with transactivation domain mode in Sp1 (Courey et al. 1989).

**Figure 5.**
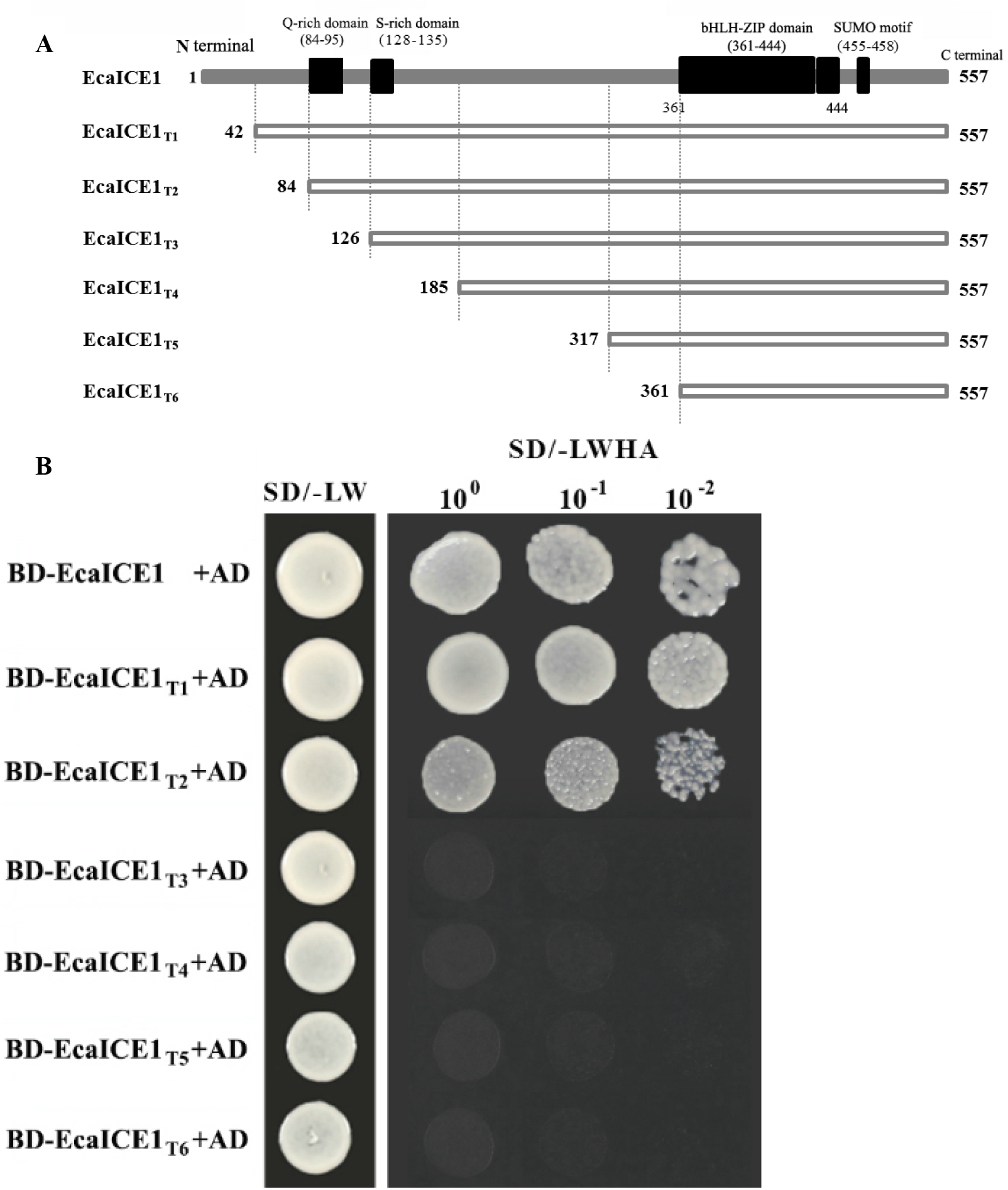
Transcriptional activation region analysis of EcaICE1 protein. BD: pGBKT7 vetor; AD: pGADT7 vetor; SD/-LW: SD/-Leu-Trp: double dropouts, SD medium with -Leu/-Trp; SD/-LWHA: quadruple dropouts, SD medium with-Leu-Trp-His-Ade;10^0^ 10^−1^ 10^−2^: dilutions with 1, 10 and 100 times respectively.

### Protein-protein interaction region between EcaICE1 with EcaSIZ1

Now the truncated EcaICE1 proteins without transcriptional activation activity (EcaICE1_T3~T6_) were fused to the GAL4 activation domain of vector pGBKT7 and co-transformed with pGADT7-EcaHOS1 into yeast strain separately to exam the protein-protein interaction region between EcaICE1 with EcaHOS1. The results showed (Figure 6) that all the truncated EcaICE1_T3_ could interact with EcaHOS1, indicating that the C-terminus protein of EcaICE1(361-557aa) is indispensable for its interaction with EcaHOS1 protein, similar to *A. thaliana* (Miura et al. 2007).

**Figure 6.**
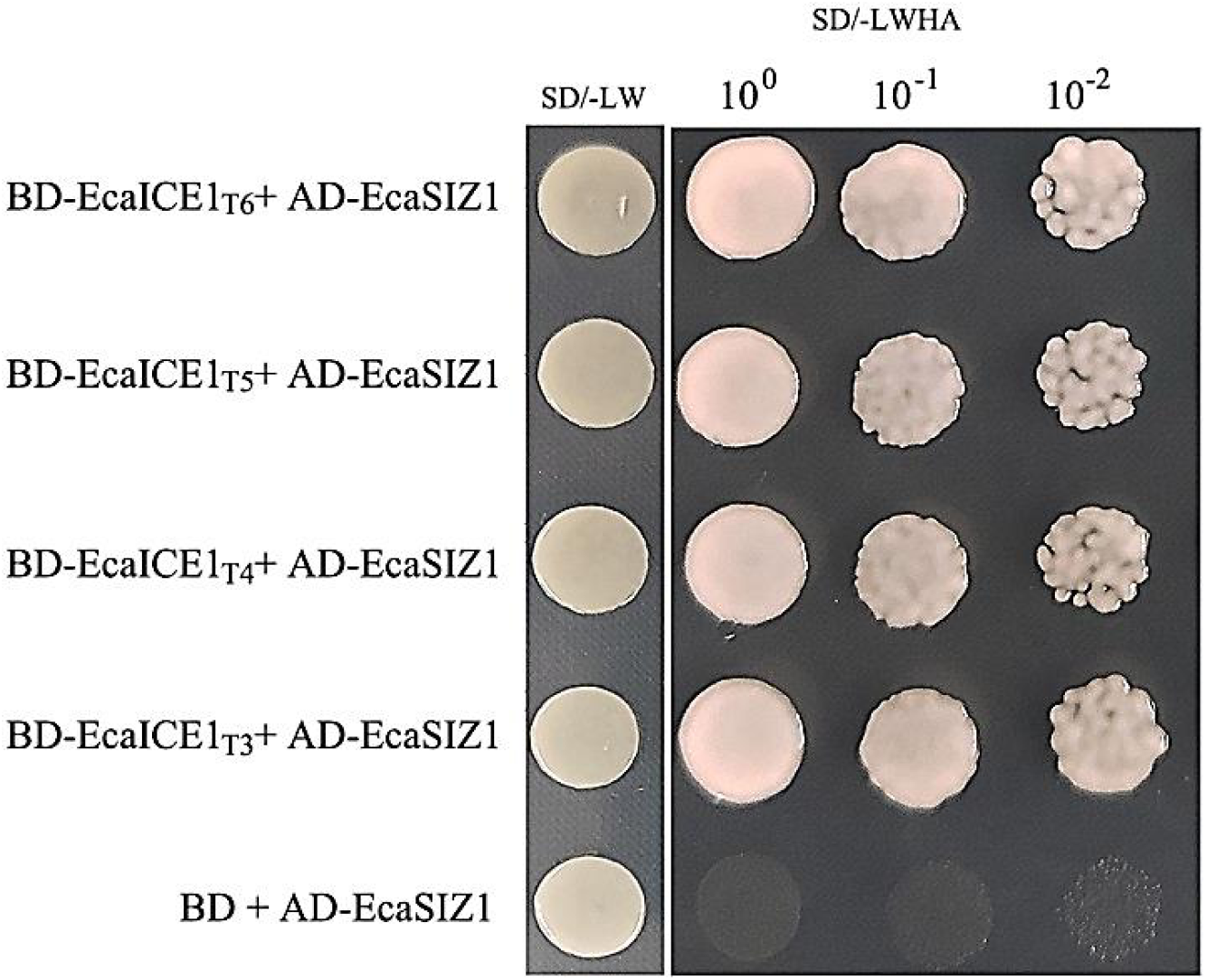
The interaction identification of EcaICE1 and EcaSIZ1 by yeast two-hybrid methods. BD: pGBKT7 vetor; AD: pGADT7 vetor; SD/-LW: SD/-Leu-Trp: double dropouts, SD medium with-Leu/-Trp; SD/-LWHA: quadruple dropouts, SD medium with-Leu-Trp-His-Ade;10^0^ 10^−1^ 10^−2^: dilutions with 1, 10 and 100 times respectively.

## DISCUSSION

It is reported that China is the world’s second largest wood consumer and the first largest wood importer (Lin 2019). At present, Chinese annual wood consumption has exceeded 600 million cubic meters, and its wood demand has become more than 50% dependent on imports. *Eucalyptus*, the main trees of man-made forest in South China even whole China, plays an important role in Chinese wooden market (Li et al. 2017). However, low temperature is a primary disadvantage factor, which causes the biomass to fall, and limits the development of its health (Janmohammadi et al. 2015). As the result, it is important to reveal the molecular mechanism of cold stress in *Eucalyptus* for improving the cold tolerance on the inheritance level as well as expanding the growing in northern region. In *Arabidopsis*, the cold signaling pathways, ICE-CBF-COR pathway (Ding et al. 2015; Shi et al. 2018) and ICE-CBF pathway is positively correlated with response to low temperature stress (Lin et al. 2014), had been studied clearly while the research of subcellular localization, transcriptional activation and interactions with upstream regulators in *E. camaldulensis* has still uncertain. Therefore, we further analyze the subcellular localization, transcriptional activation of EcaICE1, and EcaSIZ1 (the homologous gene of SIZ1) was cloned to determine the protein interactions between EcaICE1 and EcaSIZ1 as well as the key region by BiFC and yeast two-hybrid system technology for exploring the SUMOylation pathway of EcaICE1.

According to the multiple comparison of EcaICE1 protein sequence, these is a nuclear localization sequence signal in its bHLH-ZIP domain, indicated that EcaICE1 may be located in nucleus. Further, the analysis of subcellular localization demonstrates that EcaICE1 is really located in nucleus, which is similar to *P. ussuriensis* (Huang et al. 2015) and *H. brasiliensis* (Deng et al. 2017) as well as according with the characteristics of transcription factor family of plant bHLH (Gabriela et al. 2003). Above all, EcaICE1, located in nucleus, may play a regulatory role in the molecular mechanism of cold stress. What is more, we further found that the EcaICE1 84th to 95th amino acid of N-terminal, which is the Glutamine-rich domain, was identical with its homologous gene of ICE1 in *E. grandis* and *E. globulus*, and this domain only was found in the ICE1 homologous protein in *Eucalyptus*. Interestingly, it was found that EcaICE1 protein had strong transcriptional activation activity. Similar results were also reported on the ICE1 in *P. ussuriensis* (Huang et al. 2015) and *H. brasiliensis* (Deng et al. 2017) while ICE1 protein did not have transcriptional activation activity in *Arabidopsis*. The 46th to 95th and the 1th to 402th amino acid of N-terminal of protein were verified that those are the key region of transcriptional activation activity in PuICE1 of *P. ussuriensis* (Huang et al. 2015) and HbICE1 of *H. brasiliensis* (Deng et al. 2017), respectively. Unfortunately, neither of them further explored the structural characteristics of the transcriptional activation region. This study indicated that 84th to 125th amino acid of N-terminal of protein was the critical region of transcriptional activation in EcaICE1 of *E. camaldulensis*, including the Q-rich domain as well as according with the transcriptional activation structure of classical transcription factor Sp1 (Courey et al. 1989). It means that Q-rich domain may be the key transcriptional activation region of EcaICE1 protein.

SUMOylation is a posttranslational regulatory process in eukaryotes, including cold tolerance (Miura et al. 2007), ABA response (Miura et al. 2009), flowering time (Bo et al. 2010) and drought tolerance (Miura et al. 2013) and most of them are mainly devoting to *A. thaliana*. Studies in *A. thaliana* have shown that the SUMOylation pathway is mainly mediated by the SUMO E3 ligase SIZ1. Currently, SIZ1 genes have been isolated from plants such as rice (PARK et al. 2010), *Dendrobium nobile* (Liu et al. 2015), apple (Zhang et al. 2016), soybean (Cai et al. 2017) and tomato (Zhang et al. 2017), but no study on cloning the gene SIZ1 and its interaction with ICE1 in forest trees. The coding sequence of *EcaSIZ1*, which was isolated in this study, is similar to the cloned SIZ1 protein in the above plants, and also has similar protein secondary domains such as MIZ1/SP zinc finger domain, SAP domain, PINIT domain, SXS domain and PHD domain. Among them, MIZ1/SP zinc finger domain is associated with SUMO E3 ligase activity, and The SXS domain is associated with binding of SUMO (Miura et al. 2007). The SAP domain promotes binding to DNA, PINIT domain is related to intracellular retention. The above plant SIZ1 proteins are located in the nucleus and EcaSIZ1 protein should also have this conserved domain, so it indicated that EcaSIZ1 may be located in the nucleus too. PHD domain is a plant homologous domain, which only exists in SIZ1 proteins in plant (Park et al. 2010). Meanwhile, EcaSIZ1 protein shared more than 70% identity with other woody plants at the SIZ1 homologous protein level, indicating that *EcaSIZ1* is the SIZ1 gene from *E. camaldulensis*, and its encoded protein may have SUMO E3 ligase activity and mediate SUMOylation in the nucleus.

BiFC assay showed that EcaICE1 and EcaSIZ1 had the protein interaction in plant nucleus, but which region of EcaICE1 was the key region of protein interaction is still unknown. Therefore, using EcaICE1 with different truncated lengths without transcriptional activation activity as decoy protein, the 361th to 557th amino acid region at the C-terminal of EcaICE1 protein where was the key region of its interaction with EcaSIZ1 was further identified by yeast hybrid technology. The protein sequence of EcaICE1 in this region contained a SUMO domain, which was completely consistent with the result of *A. thaliana*, suggesting that the process of ICE1 SUMOylation, mediated by SIZ1, might be similar to *A. thaliana* (Miura et al. 2007). As for whether EcaSIZ1 has SUMO E3 ligase activity and whether it could couple with EcaICE1 forming the SUMO coupling for playing a role in cold toleration in plant, need further experiments to confirm.

## CONCLUSIONS

In this study, we cloned the gene *EcaICE1* and *EcaSIZ1* from *E. camaldulensis*, and the amplification of *EcaICE1* was the same as the previous result, and *EcaSIZ1* was highly conserved with other plant SIZ1 genes. EcaICE1 was similar to other ICE1 in plant, located in nucleus, and it had a strong transcriptional activation activity. Moreover, the 84-126 amino acid regions of N-terminal domain of EcaICE1 protein, were the key region of transcriptional activation activity of EcaICE1. In addition, the C-terminal region from position 361 to 557 in EcaICE1 was the key region for its interaction with EcaSIZ1. According to this study, we made a foundation on SUMOylation pathway and the molecular regulation mechanism of EcaICE1.

## Acknowledgements

This work has been supported by NSFC project 31470673. We also wish to express our appreciation to the anonymous reviewers and technical editors of the Journal of Trees for their comments and corrections of the article.

